## Supplemental information for "Risk-free polio vaccine: Recombinant expression systems for production of stabilised virus-like particles"

#### Supplemental Tables

| | PV1 SC6b (Yeast) | | PV1 SC6b (Yeast) | PV1 SC6b (Mammalian) | PV2 SC6b (Mammalian) | PV2 SC6b (Insect) |
|---|---|---|---|---|---|---|
| | DAg | CAg | GPP3 + GSH | CAg | DAg | DAg |
| | EMD-50064 | EMD-50066 | EMD-50176 | EMD-50112 | EMD-50189 | EMD-50199 |
| **Data Collection** | | | | | | |
| Microscope | Titan Krios G3i (OPIC) | | Titan Krios G3i (OPIC) | Titan Krios G3i (OPIC) | Titan Krios IV (eBIC) | Titan Krios IV (eBIC) |
| Voltage (kV) | 300 | | 300 | 300 | 300 | 300 |
| Detector | Gatan K2 | | Falcon III | Falcon III | Gatan K2 | Gatan K3 |
| Recording mode | Counting | | Linear | Linear | Counting | Super-resolution |
| Magnification (×) | 47619 | | 129629 | 129629 | 47393 | 47170 |
| Pixel size (Å) (super-resolution) | 1.05 | | 1.08 | 1.08 | 1.055 | 1.06 (0.53) |
| Defocus range (µm) | -2.9 to -0.8 | | -2.9 to -0.8 | -2.9 to -0.8 | -2.9 to -0.8 | -2.3 to -0.8 |
| Dose rate ($e^-$/pixel/s) | 7.56 | | 53.76 | 31.03 | 4.35 | 14.06 |
| Frames per movie | 30 | | 30 | 25 | 40 | 50 |
| Movie exposure time (s) | 6.00 | | 0.77 | 1.27 | 10.00 | 2.80 |
| Total electron dose ($e^-$/Å$^2$) | 41.14 | | 35.49 | 33.78 | 39.08 | 35.04 |
| **Data processing** | | | | | | |
| Movies | 4282 | | 4467 | 5104 | 1331 | 9379 |
| Initial particles (no.) | 4578 | | 159830 | 7185 | 18864 | 11032 |
| Final particles (no.) | 1320 | 2428 | 23721 | 6252 | 18378 | 3149 |
| Box size (pixels) | 400 | 400 | 448 | 400 | 480 | 400 |
| Symmetry | I1 | I1 | I1 | I1 | I1 | I1 |
| Map Resolution (Å) | 3.3 | 3.3 | 2.8 | 3.0 | 2.3 | 2.6 |
| Map sharpening *B*-factor (Å$^2$) | -61.8 | -76.9 | -146.7 | -94.0 | -67.7 | -52.2 |

Table S1: Cryo-EM data collection and image processing statistics.

| | PV1 SC6b (Yeast) | | PV1 SC6b (Yeast) | PV1 SC6b (Mammalian) | PV2 SC6b (Mammalian) | PV2 SC6b (Insect) |
|---|---|---|---|---|---|---|
| | DAg | CAg | GPP3 + GSH | CAg | DAg | DAg |
| | PDB 9EYY | PDB 9EZ0 | PDB 9F3Q | PDB 9F0K | PDB 9F59 | PDB 9F5P |
| **Model composition** | | | | | | |
| Non-hydrogen atoms | 6222 | 5090 | 6327 | 5402 | 6188 | 6184 |
| Protein residues | 794 | 654 | 804 | 685 | 772 | 790 |
| Ligands | PLM: 1 | | GPP3: 1 | | SPH: 1 | SPH: 1 |
| | | | GSH: 1 | | | |
| Waters | | | | | 123 | |
| **Refinement** | | | | | | |
| Resolution (Å) | 3.3 | 3.3 | 2.8 | 3.0 | 2.3 | 2.6 |
| Map CC[a] (Mask) | 0.87 | 0.85 | 0.89 | 0.86 | 0.89 | 0.89 |
| Map CC[a] (Volume) | 0.86 | 0.83 | 0.86 | 0.84 | 0.87 | 0.87 |
| Mean CC[a] (Ligands) | 0.78 | | 0.83 | | 0.78 | 0.80 |
| **RMS deviations** | | | | | | |
| Bond lengths (Å) | 0.003 | 0.004 | 0.003 | 0.005 | 0.003 | 0.005 |
| Bond angles (°) | 0.514 | 0.540 | 0.544 | 0.545 | 0.513 | 0.573 |
| **Mean B-factor (Å$^2$)** | | | | | | |
| Protein | 62.86 | 69.85 | 26.95 | 53.33 | 24.52 | 28.48 |
| Ligand | 58.97 | | 25.90 | | 24.41 | 26.66 |
| Water | | | | | 23.00 | |
| **Validation** | | | | | | |
| Molprobity[b] score (percentile) | 1.69 (90th) | 1.86 (83rd) | 0.98 (100th) | 1.79 (86th) | 1.08 (100th) | 1.43 (97th) |
| Clashscore[b], all atoms (percentile) | 6.18 (90th) | 6.65 (88th) | 1.68 (99th) | 3.75 (96th) | 2.25 (99th) | 4.58 (95th) |
| Ramachandran favoured (%) | 95.03 | 92.37 | 97.73 | 93.53 | 97.63 | 96.8 |
| Ramachandran allowed (%) | 4.97 | 7.47 | 2.27 | 6.47 | 2.37 | 3.2 |
| Ramachandran outliers (%) | 0.00 | 0.16 | 0.00 | 0.00 | 0.00 | 0.00 |
| Rotamer favoured (outliers) (%) | 94.79 (1.01) | 95.75 (1.06) | 94.70 (0.72) | 94.48 (1.84) | 97.58 (0.30) | 95.85 (0.89) |
| Cβ deviations >0.25 Å (%) | 0.0 | 0.0 | 0.0 | 0.0 | 0.0 | 0.0 |
| CaBLAM outliers (%) | 1.7 | 4.2 | 1.9 | 2.9 | 1.1 | 1.7 |
| CA Geometry outliers (%) | 0.52 | 0.65 | 0.13 | 0.93 | 0.40 | 0.13 |
| **EMRinger[c] score** | 4.54 | 3.68 | 5.13 | 3.80 | 6.17 | 4.84 |

Table S2: Structure refinement and validation for the capsid protein (VP0, VP1, VP3).

[a]Map CC and Mean CC (Ligands) is given for the full icosahedral virus-like particle reconstruction.
[b]Williams *et al.* (2018) Protein Sci 27:293-315.
[c]Barad *et al.* (2015) Nature Methods 12:943–946.

### Supplemental figures

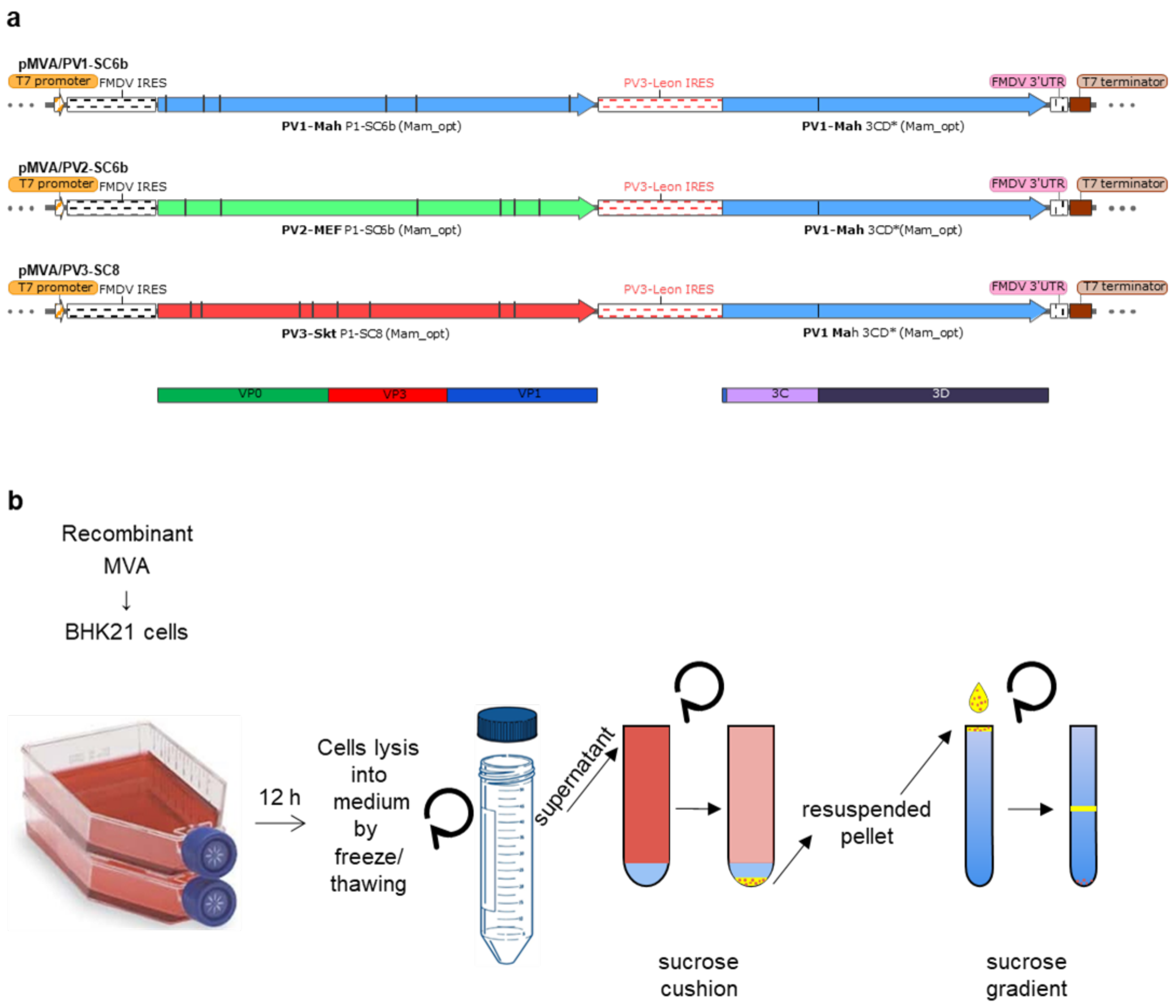


Supplemental Figure 1. Mammalian expression system

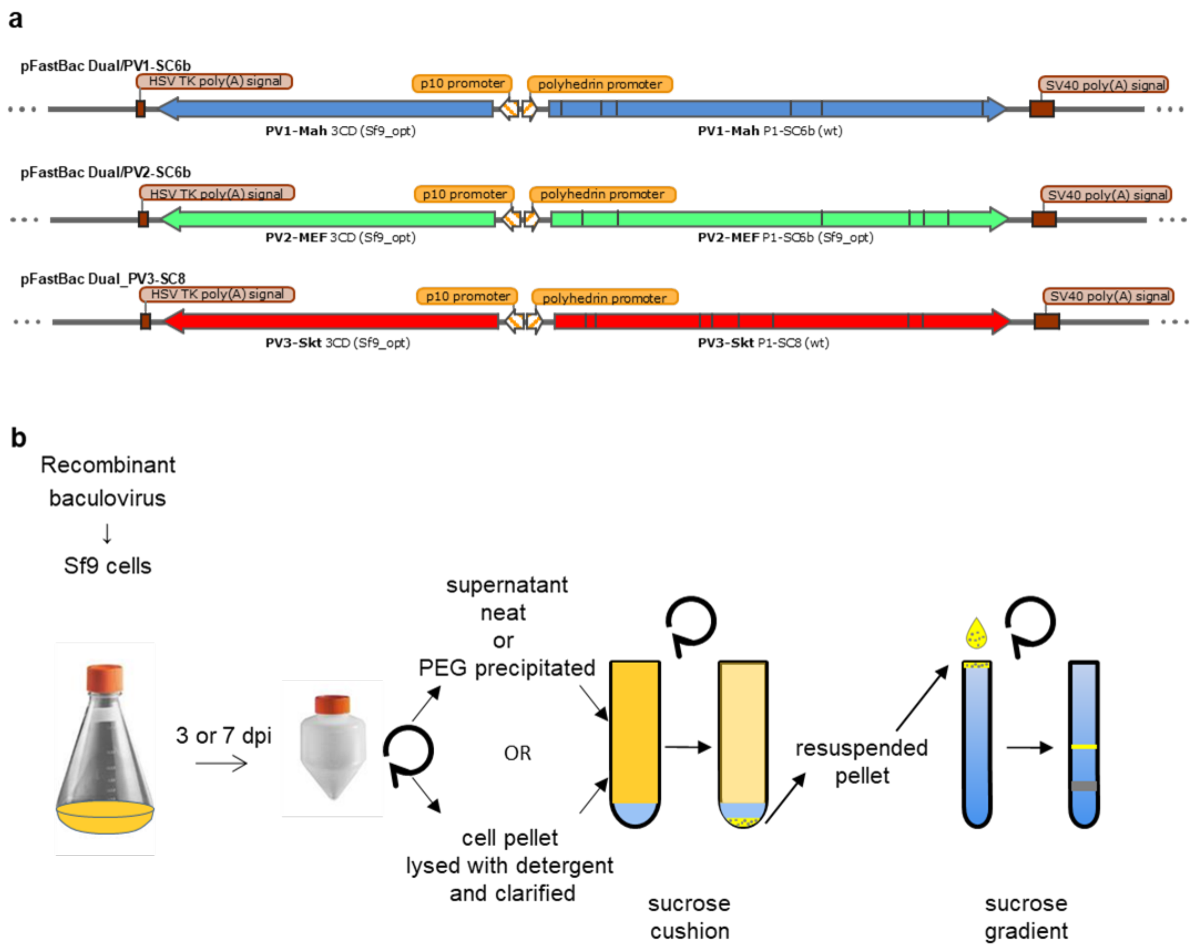


Supplemental Figure 2. Baculovirus expression system

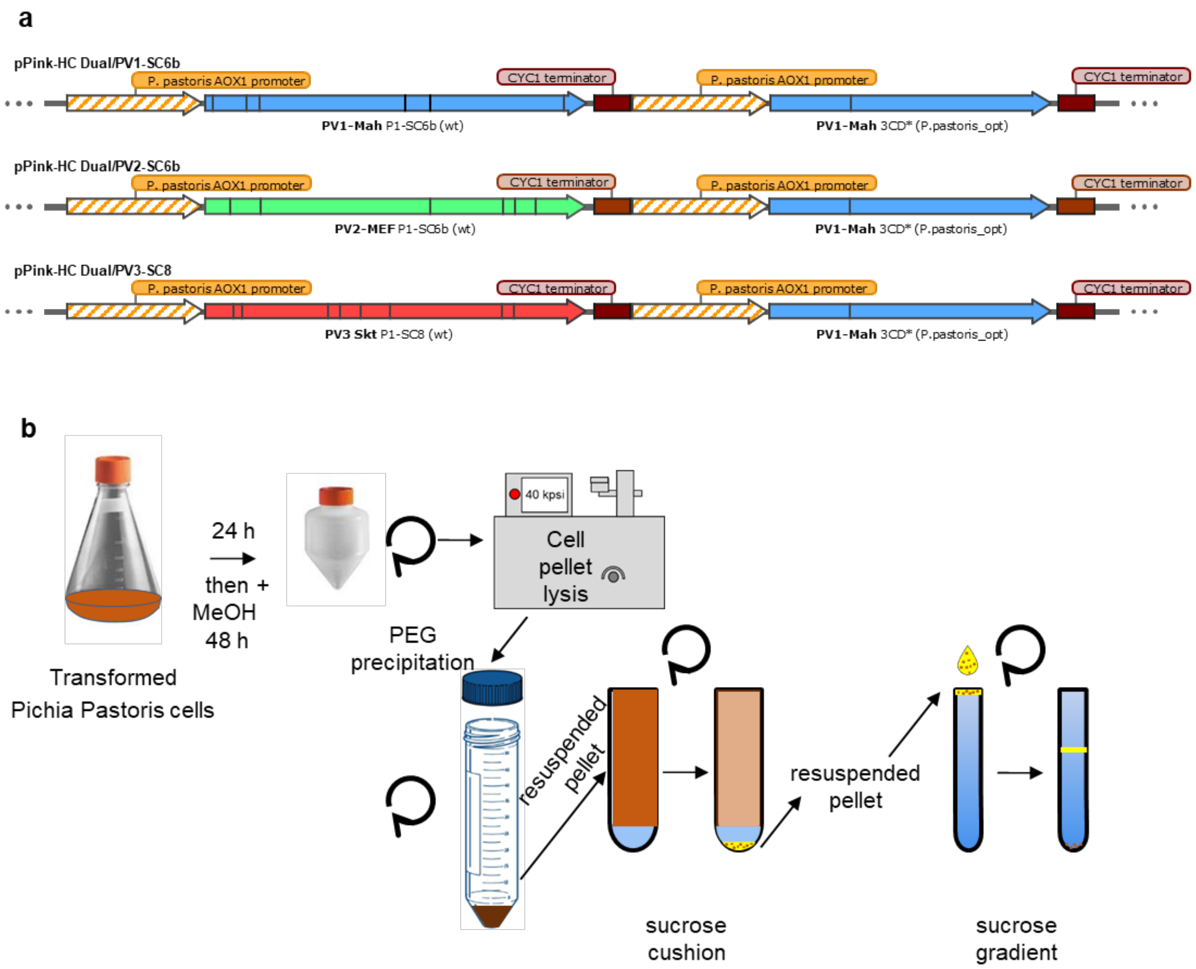


Supplemental Figure 3. Yeast expression system

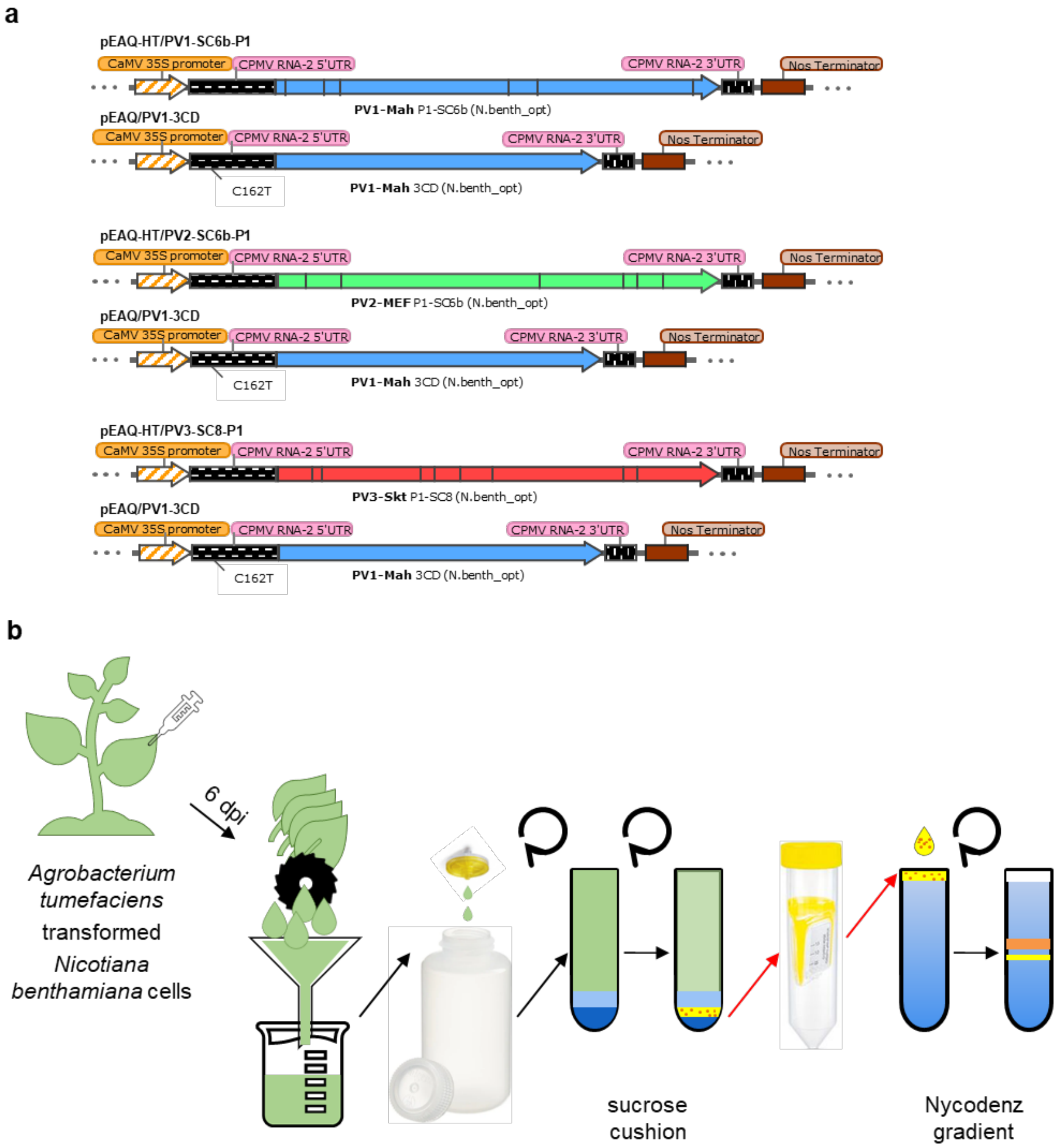


Supplemental Figure 4. Plant expression system

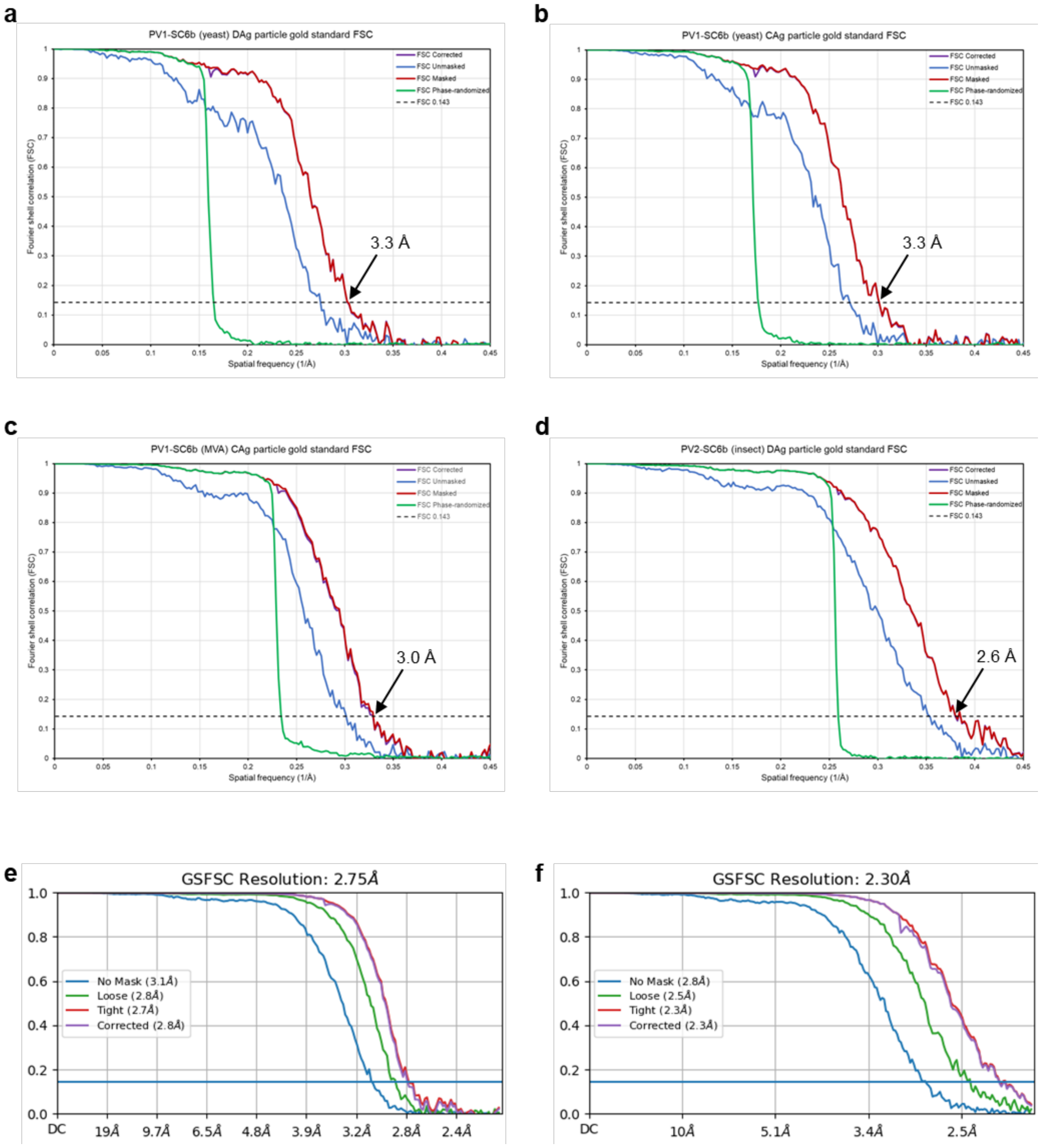


Supplemental Figure 5. Fourier Shell Correlation (FSC) resolution analysis for rsVLP cryoEM reconstructions.

**Supplemental figure legends**

Supplemental Figure 1: **Mammalian expression system**. (a) Each pMVA/PV transfer vector is transfected into primary chicken embryo fibroblasts (CEF) infected with parental MVA for homologous recombination. Mam_opt: codon optimisation for mammalian cells. Bars in the coding sequences of P1 indicate the position of the SC mutations and a bar in 3CD indicates the mutation at the junction between 3C and 3D that results in uncleavable 3CD*. (b) Mammalian cell line BHK-21 is dually infected with each MVA-PV virus alongside MVA-T7 to induce T7 promoter-mediated expression of the rsVLP cassette, followed by downstream purification from the culture supernatant into which the cells content has been released through lysis.

Supplemental Figure 2: **Insect expression system**. Each pFastBac Dual/PV transfer vector is recombined into baculovirus shuttle vector, bMON14272, resident in DH10Bac *E. coli* and the resulting recombinant baculoviruses are amplified and expressed in insect cell line Sf9. wt: native viral sequence, Sf9_opt: codon optimisation for Sf9 insect cells. Bars in the coding sequences of P1 indicate the position of the SC mutations in P1. Between 3 and 7 dpi, cultures are centrifuged and rsVLPs purified separately from the resulting cell pellet and supernatant.

Supplemental Figure 3: **Yeast expression system**. (a) Each pPink-HC-Dual/PV vector is transfected into yeast strain PichiaPink™ (selection through *ADE2* gene complementation) and expression of the PV genes controlled by *AOX1* promoters is induced by methanol. wt: native viral sequence, P. pastoris_opt: codon optimisation for *Pichia pastoris*. Bars in the coding sequences of P1 indicate the position of the SC mutations and a bar in 3CD indicates the mutation at the junction between 3C and 3D that results in uncleavable 3CD*. (b) Highly expressing PV VLP *Pichia pastoris* clones are grown to high density for 24 h in YPD medium prior to induction through the addition of methanol containing medium, YPM for 48 h when cell pellets are collected by centrifugation, followed by downstream purification.

Supplemental Figure 4: **Plant expression system**. pEAQ-HT/PVx-P1 and pEAQ/PV1-3CD plasmids are independently transformed into *Agrobacterium tumefaciens* strain LBA4404 and after amplification each pair of recombinant bacteria is mixed prior to infiltration into the leaves of *Nicotiana benthamiana* plants. N.benth_opt: codon optimisation for *Nicotiana benthamiana* plant cells. Bars in the coding sequences of P1 indicate the position of the SC mutations. The leaves are harvested 6 dpi and homogenised in a blender for rsVLP purification from the resulting sap.

Supplemental Figure 5: **Resolution analysis of PV rsVLP cryoEM reconstructions**. a-d Fourier shell correlation (FSC) calculated between two independent half sets of data as a function of spatial frequency is plotted for the (a) PV1-SC6b D Ag particle (yeast), (b) PV1-SC6b C Ag particle (yeast), (c) PV1-SC6b C Ag particle (MVA) and (d) PV2-SC6b D Ag particle (baculovirus) reconstructions from RELION. FSC is plotted for the original unmasked half-maps (blue) and masked half-maps that had density corresponding to solvent removed (red). FSC is also shown for phase-randomized half-maps (green) used to compensate for possible effects of the masking procedure before calculating the final corrected FSC (purple). Good agreement between the masked and corrected curves indicated no adverse effects from the masking. The resolution at which the corrected curve drops below the FSC=0.143 threshold (black dashed line) is indicated with an arrow. e,f Gold standard FSC curves for the (e) PV1-SC6b$^{GPP3+GSH}$ and (f) PV2-SC6b (MVA) reconstructions, respectively, generated using cryoSPARC.
